# Epstein-Barr virus co-opts ADAR1 to evade PKR activation of the integrated stress response in nasopharyngeal carcinoma

**DOI:** 10.64898/2026.10.07.757459

**Authors:** Shane Wiebe, Ming Guan Ng, Adlina Roslan, Sok Ching Cheong, Annie Wai Yeeng Chai

## Abstract

Viruses generate immunogenic double-stranded RNA (dsRNA) during their life cycle, necessitating mechanisms to evade host innate immune detection. Protein kinase R (PKR) is a pattern recognition receptor (PRR) that senses viral dsRNA and activates the host integrated stress response (ISR) by phosphorylating eukaryotic initiation factor 2 (eIF2) on its alpha subunit, causing global mRNA translation arrest. Viruses evade activation of the ISR and translation shut-down to facilitate replication. Adenosine deaminase acting on RNA 1 (ADAR1) is uniquely positioned to support viral replication due to its ability to convert adenosine (A) into inosine (I) inside dsRNA, thereby disrupting base-pair matching and preventing PKR activation. Here, we discovered that Epstein-Barr virus (EBV), the oncogenic driver of nasopharyngeal carcinoma (NPC), requires ADAR1 both to maintain latent infection and for lytic reactivation by preventing PKR-mediated ISR activation. EBV lytic reactivation stimulated viral protein expression, which was attenuated in the absence of ADAR1. Importantly, lytic reactivation or treatment of synthetic dsRNA in the absence of ADAR1 induced phosphorylation of eIF2α by PKR, stress granule formation, and caused cancer cell lethality without the production of virion progeny. Together these findings support a strategy for EBV lytic induction therapy in NPC without viral dissemination by exposing EBV to endogenous PRRs through ADAR1 inhibition.

## Introduction

The production of immunogenic double-stranded (ds)RNA creates a fundamental challenge for viruses: evading host innate immune detection. Most viruses produce dsRNA as an integral part of their life cycle [1, 2]. dsRNA is a potent activator of the integrated stress response (ISR) when detected by protein kinase R (PKR), a pattern recognition receptor (PRR) and kinase of eukaryotic initiation factor 2 (eIF2; a heterotrimer comprised of the α, β, and γ subunits). Phosphorylation of eIF2 on the alpha subunit rapidly inhibits global translation and paradoxically upregulates stress-response genes to preserve the cell under stress conditions. Mechanistically, p-eIF2α inhibits eIF2B, a guanine nucleotide exchange factor (GEF), from recycling GDP to GTP [3–5]. GTP is an integral component of the ternary complex, and together with eIF2 and the methionine transfer RNA (Met-tRNAi), is required for the 40S ribosomal subunit to begin scanning and initiate translation [6]. Viruses require mechanisms that evade PKR and activation of the ISR to allow translation of their mRNAs [7].

Adenosine deaminase acting on RNA 1 (ADAR1) is an RNA-editing enzyme that converts adenosine (A) into inosine (I) inside dsRNA. Since inosine resembles guanosine, this creates a base pair mismatch and disruption of the dsRNA structure [8, 9]. A-to-I editing is critical to prevent aberrant activation of the ISR by PKR sensing of endogenous dsRNA, such as inverted Alu repeat elements within pre-mRNAs and microRNA precursors [10]. However, this function situates ADAR1 as an ideal host factor to be hijacked by viruses to evade innate immune detection of their dsRNA by PKR. Moreover, ADAR1 can directly bind and inhibit PKR through its dsRNA binding domain 3 (dsRBD3) [11].

The Epstein-Barr virus (EBV)-encoded EBNA1 protein was recently shown to induce translation of ADAR1 mRNA by forming a trimeric complex with insulin-like growth factor 2 mRNA-binding protein 3 (IGF2BP3) and eukaryotic translation initiation factor 4G1 (eIF4G1) [12]. This was the first evidence that EBV actively maintains ADAR1 expression and suggested that ADAR1 may have an important role in EBV’s life cycle. In addition, Zhang et al. showed that ADAR1 is required for productive lytic reactivation from latency of another gammaherpesvirus, Kaposi’s sarcoma-associated herpesvirus (KSHV), by attenuating interferon (IFN) responses [13].

Given these findings, we hypothesized that EBV requires ADAR1 to prevent PKR-induced activation of the ISR and translational shut-down to maintain production of viral lytic proteins during lytic reactivation. To this end, we employed several nasopharyngeal carcinoma (NPC) cell line models, since persistent latent EBV infection drives NPC oncogenesis [14], while retaining the capacity to be induced into lytic reactivation. NPC also has some of the highest rates of A-to-I editing among cancers, suggesting active ADAR activity [15].

Our study reveals that NPCs are highly dependent on ADAR1 for survival and to prevent PKR phosphorylation, even in the absence of EBV. We demonstrate that ADAR1 is required for EBV persistent latent infection and lytic reactivation by preventing PKR-induced ISR activation and stress granule formation. Lytic reactivation in the absence of ADAR1 abolished productive viral replication while simultaneously inducing death of the cancerous host cell. We propose that exposing EBV to the ISR by inhibiting ADAR1 will enhance the efficacy of lytic induction therapy in NPC while precluding viral dissemination, thereby solving a major limitation of the current cytolytic treatment strategy in EBV-driven malignancies.

## Results

### NPC cells are dependent on ADAR1 for survival and to prevent PKR activation

We first assessed whether NPC cells exhibit any dependency on ADAR1 alone. One of the features of NPC is high levels of dsRNA editing [15]. ADAR1 is a well-established oncogene in several cancers, including head and neck cancer [16]. Recent reports have indicated that NPC patients with high ADAR1 expression have worse survival [12]. We therefore hypothesized that NPCs are dependent on ADAR1. We compared ADAR1 dependency in both EBV-positive and EBV-negative NPCs.

Western blot analysis confirmed successful knockdown of ADAR1 using two sgRNAs (sgA1K and sgA2G) in HK-1 Cas9 (EBV-negative, Fig. 1A, left panel); HK-1 EBV Cas9 (EBV-positive, Fig. 1B, left panel); C666-1 Cas9 (EBV-positive, Fig. 1C, left panel); and NPC268 Cas9 (EBV-positive, Fig. 1D, left panel) compared to their respective sgNT (non-targeting) control. Given the important role of ADAR1 in mitigating innate immune responses by dsRNA sensors, we then investigated whether the major dsRNA detection pathways, namely PKR and Melanoma Differentiation-Associated protein 5 (MDA5), were activated upon loss of ADAR1. In the sgA1K and sgA2G groups, we observed increased phosphorylation of the dsRNA sensor PKR relative to total PKR, but no change in MDA5 protein levels (Fig. 1A-D, left panels). Since MDA5 expression is induced in response to inflammatory agents such as dsRNA [17], MDA5 protein levels serve as a supporting marker of MDA5 pathway activity. These findings therefore suggest that ADAR1 preferentially inhibits dsRNA sensing by PKR, rather than MDA5, in NPC.

**Figure 1:**
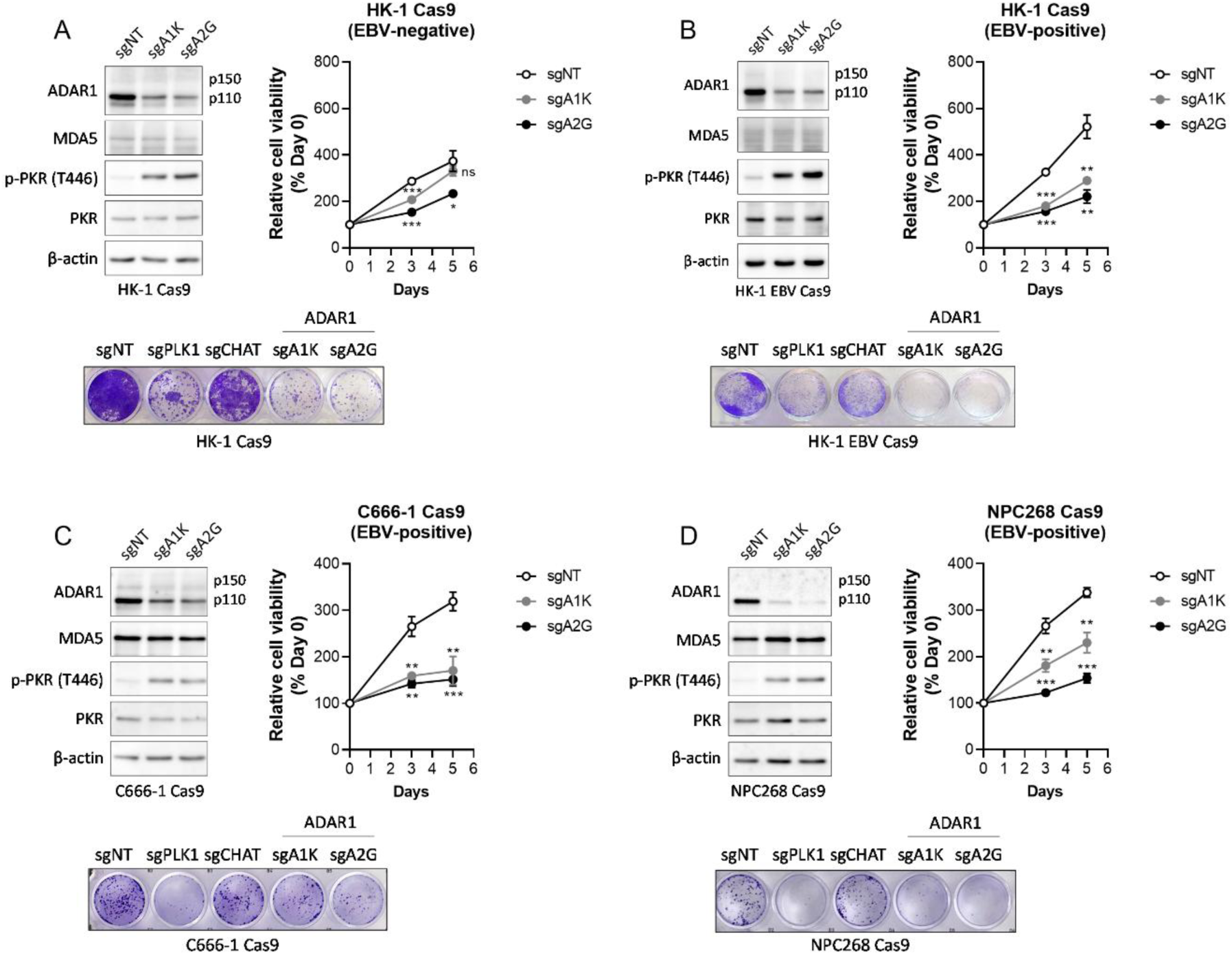
NPC cells are dependent on ADAR1 for survival and to prevent PKR activation. **A** Analysis of ADAR1 dependency in HK-1 Cas9 (EBV-negative). Left panel: Western blot validation of ADAR1 knockdown and analysis of PKR activation and MDA5 protein abundance. β-actin was used as housekeeping. Right panel: MTT analysis of cell viability over five days in sgNT vs. sgA1K or sgA2G. Each group was normalized to their respective day 0 MTT measurement. Bottom panel: Colony formation analysis by crystal violet staining in sgNT vs. sgA1K or sgA2G. sgPLK1 was included as a dependent positive control; sgCHAT was a non-dependent negative control. The western blot, MTT and CFA analysis were similarly conducted in **B** HK-1 Cas9 (EBV-positive), **C** C666-1 Cas9 (EBV-positive), and **D** NPC268 Cas9 (EBV-positive) NPC cell lines. N=6 per group. Western blots were reproduced in at least two independent experiments. Data are presented as mean ± s.e.m.; ***p<0.001; **p<0.01; *p<0.05; ns, not significant; calculated using a Two-way ANOVA with Dunnett’s multiple comparisons test.

MTT growth curve analysis revealed that each cell line was dependent on ADAR1 for viability, regardless of EBV status (Fig. 1A-D, right panels). We further validated ADAR1 dependency in NPC by measuring colony formation. Both sgA1K and sgA2G impaired colony formation in each NPC cell line compared to sgNT (Fig. 1A-D, bottom panels)—sgPLK1 was included as an essential gene positive control, and sgCHAT as a non-essential negative control.

Taken together, these data demonstrate that NPC cells, regardless of EBV status, are dependent on ADAR1 for viability and to prevent PKR activation.

### EBV requires ADAR1 to maintain latent gene expression, persistent latent infection and lytic reactivation in NPC cells by inhibiting PKR

EBV persistent latent infection drives oncogenesis in NPC [14]. We reasoned that ADAR1 has a role in maintaining EBV latent infection which may contribute to the NPC dependency on ADAR1. Given previous findings that EBNA1 induces ADAR1 expression [12], we first compared ADAR1 expression in isogenic NPCs based on EBV status. Consistently, we observed higher expression of both ADAR1 p150 and p110 isoforms in EBV-positive NPCs compared to their isogenic EBV-negative counterparts (Fig. 2A). Deleting ADAR1 in NPC268 Cas9 cells significantly impaired expression of EBV latent mRNAs, including EBNA1, EBER1/2, LMP1 and LMP2 (Fig. 2B).

**Figure 2:**
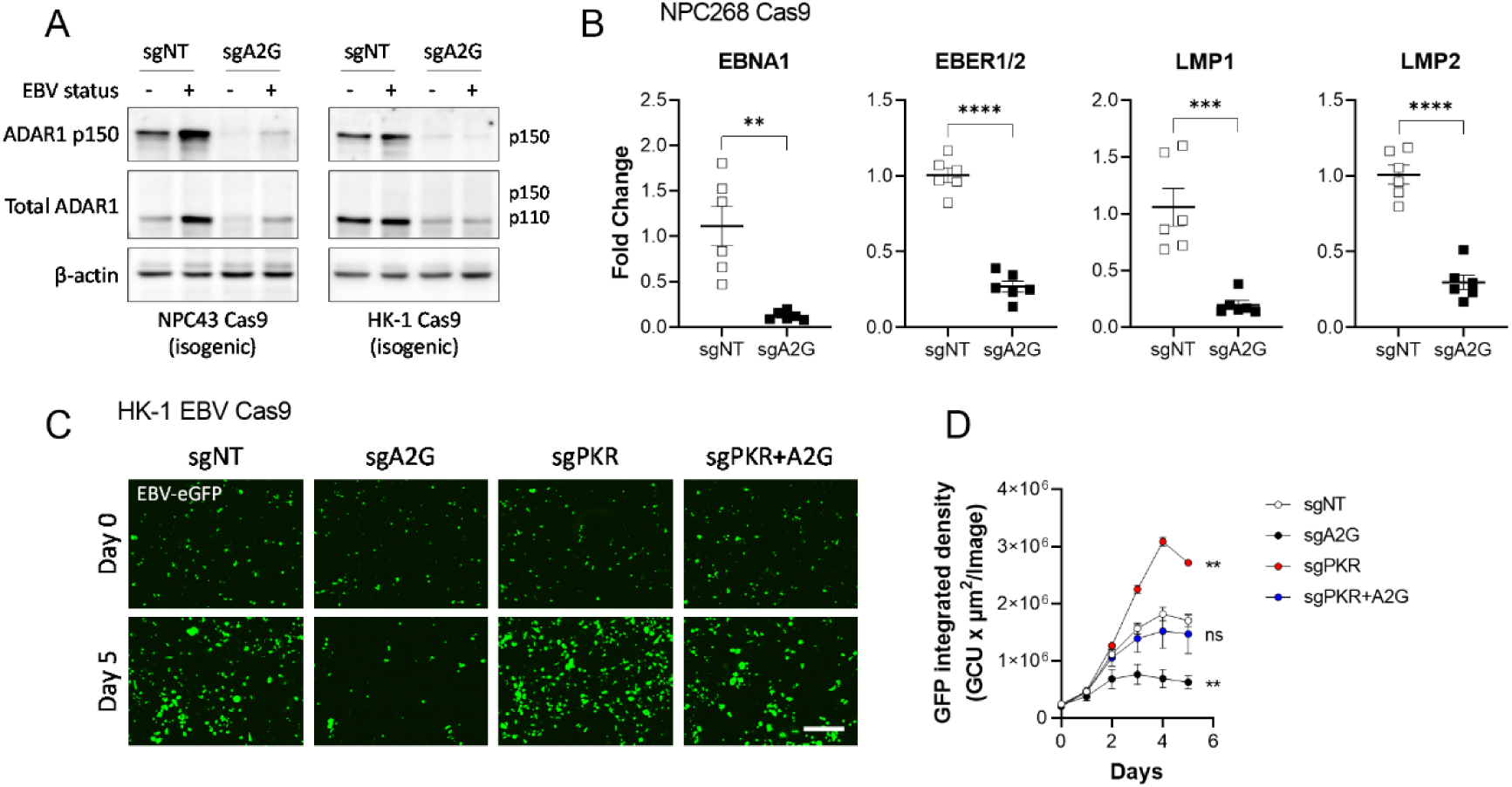
EBV requires ADAR1 to maintain latent gene expression and persistent latent infection in NPC cells by inhibiting PKR. **A** Western blot of ADAR1 protein levels in isogenic NPC43 Cas9 (left panel) and HK-1 Cas9 (right panel) comparing EBV status in sgNT and sgA2G groups. β-actin was used as housekeeping. Western blots were reproduced in at least two independent experiments. **B** qPCR analysis of EBV latent genes, EBNA1, EBER1/2, LMP1, and LMP2 in NPC268 Cas9 in sgNT vs. sgA2G. N=6 per group. **C** Incucyte fluorescent live-cell imaging of EBV-eGFP expression in HK-1 EBV Cas9 cells over five days in sgNT, sgA2G, sgPKR, and sgPKR+A2G groups. Scale bar=400 µm. **D** Quantification of EBV-eGFP expression by integrated density in sgNT, sgA2G, sgPKR, and sgPKR+A2G over five days. Images were captured every six hours and binned for each day. N=3 per group. Results were replicated in independent experiments. Data are presented as mean ± s.e.m.; ****p<0.0001; ***p<0.001; **p<0.01; *p<0.05; ns, not significant; calculated using a One-way student’s T-Test or a Two-way ANOVA with Dunnett’s multiple comparisons test.

We then assessed how ADAR1 regulates latent EBV during cell growth using live-cell imaging. HK-1 Cas9 cells were exogenously infected with a recombinant EBV expressing an eGFP reporter (referred to as HK-1 EBV Cas9), as previously reported [18]. eGFP integrated density was visualized over five days of cell growth (Fig. 2C). As cells proliferated, the eGFP signal increased in the sgNT group (Fig. 2C and D). In sgA2G, eGFP signal was attenuated compared to sgNT during cell growth, but was rescued when PKR was simultaneously deleted (sgPKR+A2G) (Fig. 2C and D).

The previous findings that ADAR1 is required for productive lytic reactivation from latency of KSHV [13] prompted us to assess EBV lytic gene expression during lytic reactivation. Deletion of ADAR1 robustly attenuated expression of lytic genes BZLF1, BRLF1, BMRF1, and BLLF1 when EBV was induced into lytic reactivation by either transfecting BZLF1 (Fig. 3A) or treating cells with TPA (12-O-tetradecanoylphorbol-13-acetate) and NaB (sodium butyrate) (Fig. 3B). Importantly, deleting ADAR1 prevented the generation of viral progeny, measured by EBV copy number, upon lytic reactivation (Fig. 3C).

**Figure 3:**
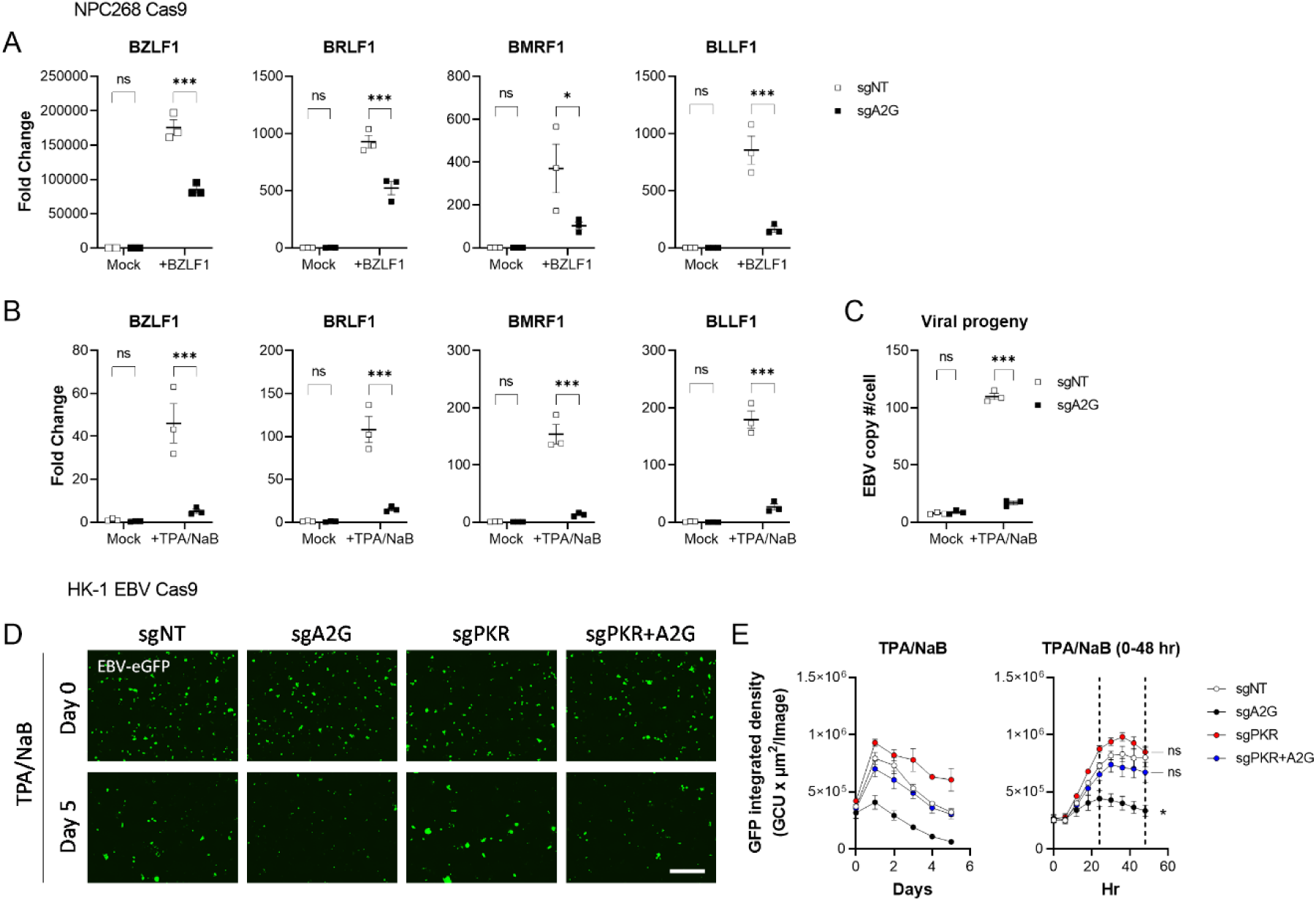
EBV requires ADAR1 for lytic reactivation and generation of viral progeny in NPC cells by inhibiting PKR. **A** qPCR analysis of EBV lytic genes, BZLF1, BRLF1, BMRF1, and BLLF1 in NPC268 Cas9 in sgNT vs. sgA2G, two days following lytic reactivation by mock or BZLF1 transfection (1 µg/mL). N=3 per group. **B** qPCR analysis of EBV lytic genes, BZLF1, BRLF1, BMRF1, and BLLF1 in NPC268 Cas9 in sgNT vs. sgA2G, two days following lytic reactivation by mock or TPA (20 ng/mL) and NaB (2 mM) treatment. N=3 per group. **C** Quantification of EBV copy number by qPCR of EBNA1 genomic DNA in NPC268 Cas9 in sgNT vs. sgA2G, two days following lytic reactivation by mock or TPA (20 ng/mL) and NaB (2 mM) treatment. N=3 per group. **D** Incucyte fluorescent live-cell imaging of EBV-eGFP expression in HK-1 EBV Cas9 cells over five days in sgNT, sgA2G, sgPKR, and sgPKR+A2G groups following lytic reactivation by TPA (20 ng/mL) and NaB (2 mM). Scale bar=400 µm. **E** Quantification of EBV-eGFP expression by integrated density in sgNT, sgA2G, sgPKR, and sgPKR+A2G over five days. Images were captured every six hours and binned for each day (left panel). The same data is displayed with six-hour time intervals to visualize eGFP expression dynamics withing the first 48 hr (right panel). N=3 per group. Results were replicated in independent experiments. Data are presented as mean ± s.e.m.; ***p<0.001; *p<0.05; ns, not significant; calculated using a Two-way ANOVA with Dunnett’s or Bonferroni’s multiple comparisons test.

We then assessed the time course dynamics of EBV lytic reactivation using HK-1 EBV Cas9 by live-cell imaging. Treating the cells with TPA/NaB induced lytic reactivation and eGFP expression in control sgNT that peaked between 24 and 48 hr and then declined steadily on the following days (Fig. 3D and E). eGFP expression was significantly attenuated upon ADAR1 deletion, but was rescued to control levels in the absence of PKR (Fig. 3D and E).

Given these data, we conclude that EBV requires ADAR1 for persistent latent infection and productive lytic reactivation through ADAR1-mediated regulation of PKR.

### EBV lytic reactivation and synthetic dsRNA activate PKR-mediated integrated stress response in the absence of ADAR1 which prevents EBV lytic protein production

We next assessed the molecular events occurring during the EBV lytic reactivation time course. By 48 hr, lytic reactivation by TPA/NaB treatment induced Zta protein expression, encoded by the BZLF1 gene, in both HK-1 EBV Cas9 and NPC268 Cas9 NPC cells (Fig. 4A and B). In the absence of ADAR1, Zta protein levels were reduced or nearly absent with a concomitant increase in p-PKR and p-eIF2α relative to their total proteins (Fig. 4A and B).

**Figure 4:**
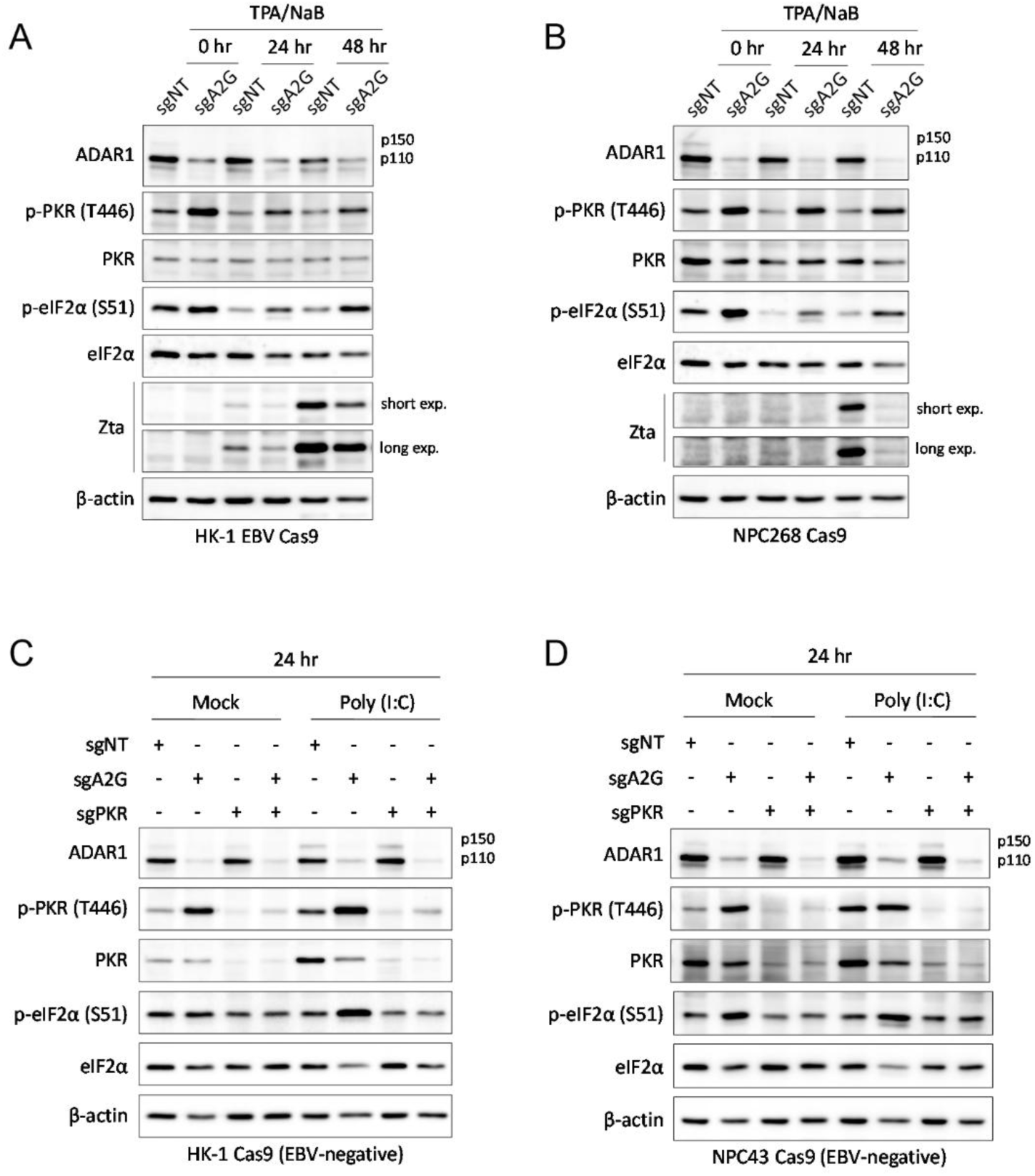
EBV lytic reactivation and synthetic dsRNA activate PKR-mediated integrated stress response in the absence of ADAR1 which prevents EBV lytic protein production. Western blot analysis of ISR activation via PKR in sgNT vs. sgA2G at 0 hr, 24 hr, and 48 hr following EBV lytic reactivation by TPA (20 ng/mL) and NaB (2 mM) treatment in **A** HK-1 EBV Cas9 and **B** NPC268 Cas9 cells. β-actin was used as housekeeping. Western blot analysis of ISR activation via PKR in sgNT, sgA2G, sgPKR and sgPKR+A2G after 24 hr of mock or poly(I:C) (10 µg/mL) transfection in **C** HK-1 Cas9 (EBV-negative) and **D** NPC43 Cas9 (EBV-negative) cells. β-actin was used as housekeeping. Western blots were reproduced in at least two independent experiments.

To further clarify the role of viral dsRNA in the activation of PKR and the integrated stress response during lytic reactivation in the absence of ADAR1, we treated EBV negative NPC cells with poly(I:C) synthetic dsRNA for 24 hr and assessed ISR activation. Consistently, poly(I:C) induced phosphorylation of PKR and eIF2α which was exaggerated in the absence of ADAR1, in both EBV-negative HK-1 Cas9 and NPC43 Cas9 cells (Fig. 4C and D). Co-deleting PKR and ADAR1 prevented activation of the ISR in response to poly(I:C) (Fig. 4C and D).

Collectively, our findings establish that ADAR1 prevents PKR-mediated activation of the ISR to allow EBV lytic protein expression and progression through the lytic cycle.

### ADAR1 prevents stress granule formation in NPC cells

Phosphorylation of eIF2α causes global translation arrest and induces stress granule (SG) formation of translationally inactive mRNAs [19]. Ras GTPase-Activating Protein-Binding Protein 1 (G3BP1) is an RNA binding protein that aggregates into SGs and serves as a reliable SG marker during ISR activation [20].

We first assessed SG formation following viral dsRNA stress using poly(I:C). Treating EBV-negative HK-1 Cas9 cells with poly(I:C) or deleting ADAR1 alone was sufficient to induce SG formation, but poly(I:C) treatment in ADAR1 knockdown led to a combined increase in SG formation (Fig. 5A). We then assessed SG formation during EBV lytic reactivation. Similar to poly(I:C) treatment, lytic reactivation induced SG formation after 48, but not 24 hrs (Fig. 5B and C). Treatment of TPA/NaB in the absence of ADAR1 resulted in further SG formation, but only at 48 hrs (Fig. 5B and C).

**Figure 5:**
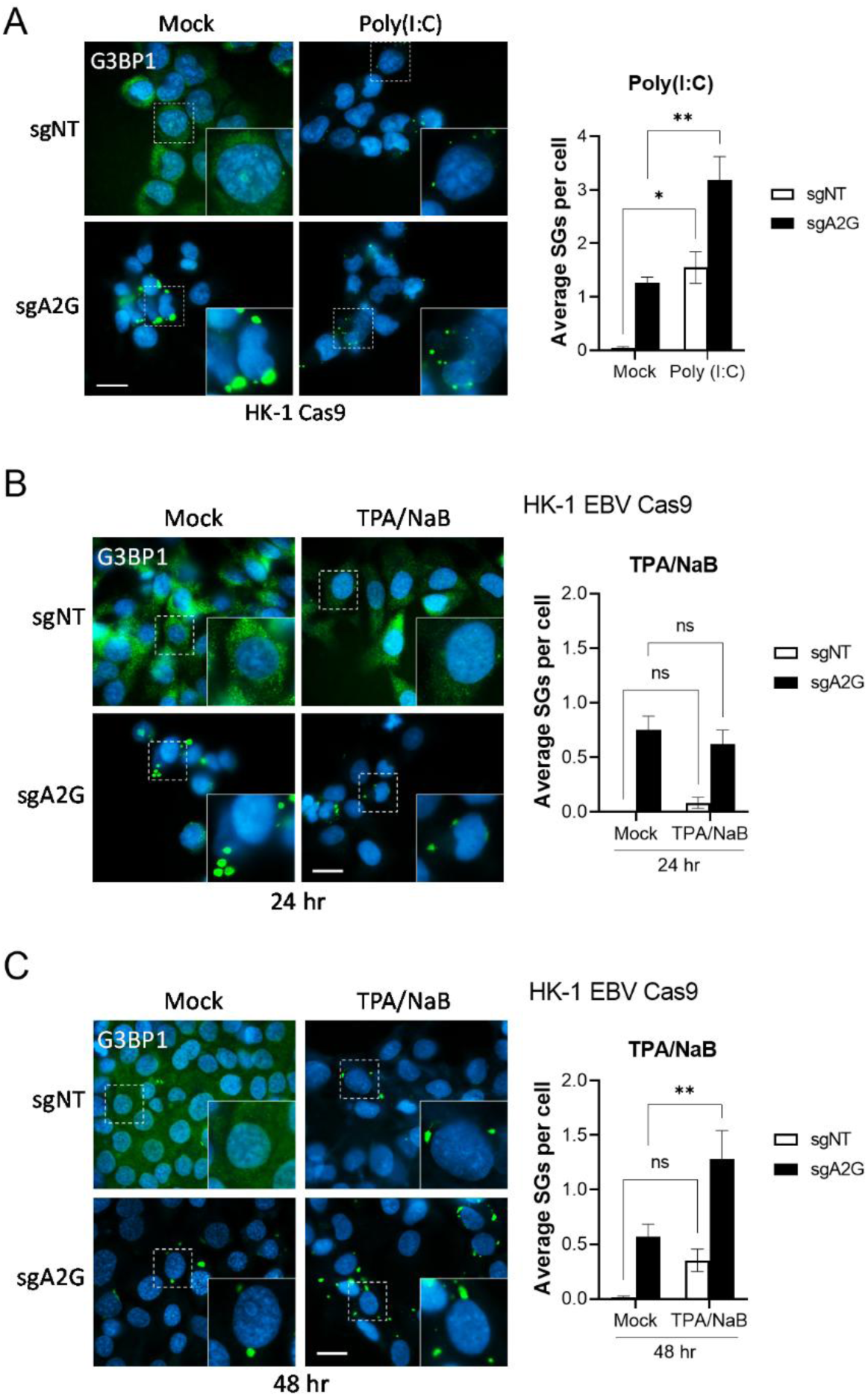
ADAR1 prevents stress granule formation in NPC cells. **A** (Left panel) Stress granules were visualized by immunofluorescence microscopy in HK-1 EBV Cas9 cells in sgNT vs. sgA2G after 6 hr of mock or poly(I:C) (1 µg/mL) transfection. Blue=Hoechst-stained nucleus; green=G3BP1. Right panel: Quantification of average stress granules per cell. N=3-5 images per group, averaged from at least 30 cells. **B-C** (Left panels) Stress granules were visualized by immunofluorescence microscopy in HK-1 EBV Cas9 cells in sgNT vs. sgA2G after **B** 24 hr or **C** 48 hr of mock or TPA (20 ng/mL) and NaB (2 mM) treatment. Blue=Hoechst-stained nucleus; green=G3BP1. (Right panels) Quantification of average stress granules per cell. N=3-5 images per group, averaged from at least 30 cells. Scalebar=20 µm. Immunofluorescence results were replicated in at least two independent experiments. Data are presented as mean ± s.e.m.; **p<0.01; *p<0.05; ns, not significant; calculated using a Two-way ANOVA with Šídák’s multiple comparisons test.

Taken together, these findings support the conclusion that EBV lytic reactivation induces SG formation, most likely through activation of the ISR by viral dsRNA, and is attenuated by ADAR1.

### Inducing EBV lytic reactivation in the absence of ADAR1 is lethal to NPC cells and constitutes a potential combination therapy

In the absence of ADAR1, PKR senses dsRNA unhindered and subsequently activates the ISR. Phosphorylation of eIF2α induces SG formation and inhibits global translation, including the translation of viral lytic mRNAs. The end result is impaired generation of viral progeny and cell death, likely due to prolonged activation of the stress response pathway. We found that although deletion of ADAR1 alone or induction of lytic reactivation alone impairs cell viability, the combination of both results in near complete loss of cell viability (Fig. 6A and B).

**Figure 6:**
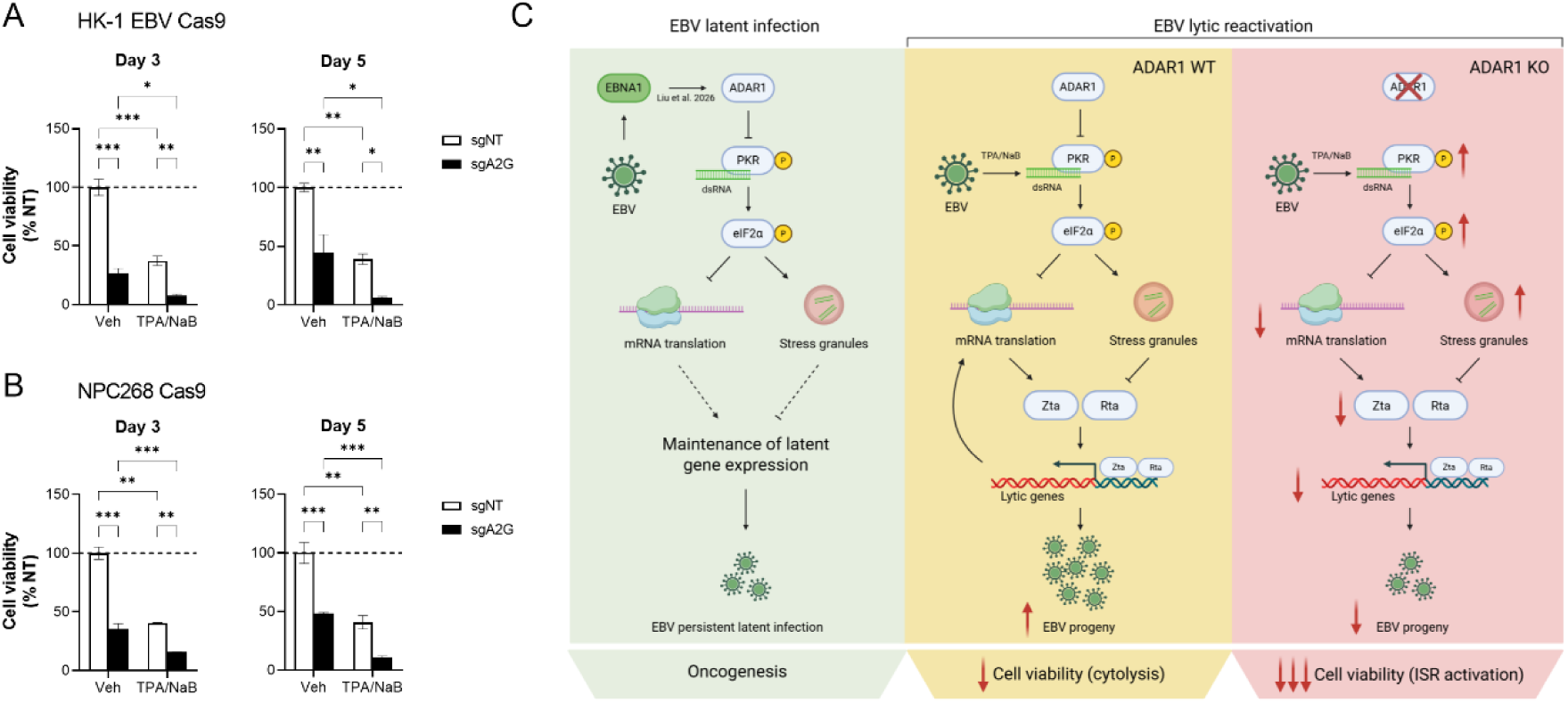
Inducing EBV lytic reactivation in the absence of ADAR1 is lethal to NPC cells and constitutes a potential combination therapy. MTT analysis of cell viability in **A** HK-1 EBV Cas9 and **B** NPC268 Cas9 cells at day 3 and day 5 in sgNT vs. sgA2G treated with either vehicle (Veh) or TPA/NaB. N=3. Data are presented as mean ± s.e.m.; ***p<0.001; **p<0.01; *p<0.05; calculated using a Two-way ANOVA with Šídák’s multiple comparisons test. **C** Working model for how EBV promotes latent infection and oncogenesis through ADAR1-mediated suppression of PKR and the integrated stress response, while leveraging this mechanism to evade cell death during lytic reactivation.

Our collective findings support a model that in order for EBV to successfully replicate during lytic reactivation, it requires ADAR1 to safeguard against dsRNA sensing-induced PKR activation (Fig. 6C). By preventing PKR activation, p-eIF2α remains at baseline levels, thereby preventing SG formation and allowing translation of EBV lytic mRNAs and generation of viral progeny.

## Discussion

In this study, we found that EBV requires ADAR1 to prevent PKR sensing of dsRNA and activation of the ISR during forced lytic reactivation and to maintain persistent latent infection. NPC cell viability was dependent on ADAR1 and the presence of EBV enhanced ADAR1 protein levels (Fig. 1 and Fig. 2A)—potentially through EBNA1-mediated translational upregulation, as previously shown [12]. Deleting ADAR1 disrupted both latent and lytic EBV mRNA expression (Fig. 2B and Fig. 3A and B). Reduced EBV mRNA expression in the absence of ADAR1 is likely a downstream consequence of ISR activation and translational shutdown of the viral proteins necessary for EBV gene transcription. For example, new EBV lytic mRNAs would not be expressed without their requisite transcription factor, Zta [21] (Fig. 4 A-B; Fig. 6C). Consistent with our findings, inhibition of the ISR by EBV latent membrane protein 1 (LMP1) was also shown to be necessary for EBV to maintain latent infection in NPC [22]. ADAR1-mediated suppression of the PKR–ISR pathway is therefore essential for EBV to maintain the viral gene expression required for latent infection and during lytic reactivation (Fig. 6C).

ADAR1 preferentially regulates dsRNA sensing through PKR, as deleting ADAR1 in NPC cells had little effect on MDA5 protein abundance (Fig. 1A-D, left panels). This is in contrast to previous reports, which demonstrated that ADAR1 prevents MDA5 activation and downstream IFN responses to support lytic reactivation of another gammaherpesvirus, KSHV [13]. However, similar to EBV, KSHV also requires PKR inhibition for efficient lytic reactivation: the KSHV protein ORF57 binds PKR and PKR-activating protein (PACT), preventing their interaction and ISR activation [23]. Although the role of ADAR1 in regulating PKR in the context of KSHV infection remains to be fully elucidated, these findings in a closely related virus further support our model that viral dsRNA produced during lytic reactivation is otherwise detrimental to gammaherpesviruses, necessitating mechanisms to suppress its recognition by PKR.

We argue that, like most viruses, cytosolic dsRNA species accumulate in NPC cells during EBV lytic reactivation [1, 2]. This increase in dsRNA is likely responsible for PKR activation in the absence of ADAR1. For instance, transfecting EBV-negative NPC cells with synthetic poly(I:C) dsRNA recapitulates the induction of PKR-ISR signalling and SG formation observed following EBV lytic reactivation, providing orthogonal support for this mechanism. Because poly(I:C) does not contain adenosines available for A-to-I editing, ADAR1 suppression of PKR is not expected to be editing-dependent. Rather, ADAR1 likely inhibits PKR through direct protein-protein interaction. Studies have demonstrated direct ADAR1-PKR interaction [24], specifically between the ADAR1 dsRBD3 and PKR kinase domain [11], which inhibits PKR activation. ADAR1-PKR binding is further enhanced during HIV-1 [25] and HSV-1 infection [26] to prevent viral-induced PKR activation. Future studies should investigate the dynamics of ADAR1-PKR interaction during EBV lytic reactivation, as well as identify critical binding residues for the development of small molecule inhibitors that disrupt ADAR1-PKR binding for use in combination with lytic inducing compounds.

In NPC, EBV drives oncogenesis in a persistent latent infection, making it ideally suited for lytic induction therapy by forcing the virus into lytic reactivation [27]. Lytic induction (or cytolytic) therapy overcomes limitations in directly targeting EBV in a latent infection, where EBV only expresses a subset of viral proteins, hence with limited potential oncogenic targets. Without active replication, EBV is also insensitive to most antiviral drugs. EBV can be induced into lytic reactivation using compounds such as histone deacetylase (HDAC) inhibitors (eg. NaB [28]), chemotherapeutics (eg. gemcitabine [29]), epigenetic modulators (eg. 5-aza-cytidine [30]), or small molecules targeting host signalling pathways (eg. the protein kinase C activator, TPA [31, 32]). When treated in combination with antiviral drugs, such as ganciclovir [33], lytic induction therapy has the potential to treat NPC and other EBV associated malignancies by selectively killing the infected cancerous host cell.

One of the major limitations of lytic induction therapy is the consequent production of new viral progeny and dissemination to neighbouring cells during treatment [34]. Moreover, only a small proportion of latently infected cells are successfully reactivated, leaving the majority untreated [35]. Several compounds induce lytic reactivation without generating infectious viral progeny, such as the bacteriostatic antibiotic—clofoctol [36]. Consistent with our findings, clofoctol prevents EBV protein expression and virion production by activating the ISR—specifically through the PKR-like endoplasmic reticulum kinase (PERK) pathway [36]. These results provide converging evidence that triggering the ISR effectively blocks productive lytic replication. We propose that exposing EBV’s dsRNA to PKR sensing by inhibiting ADAR1 during lytic reactivation can both prevent viral protein expression and cause death of the cancerous host cell. Importantly, the ISR will only be activated in the EBV-positive cancerous cells during lytic induction therapy, thereby limiting damage to healthy tissue. Inhibiting ADAR1 may therefore be a critical component for lytic induction therapy to prevent viral dissemination.

In recent years, several ADAR1 inhibitors have been developed, including the spliceosome-modulating ADAR1 p150 inhibitor rebecsinib [37]; the Zα-domain-targeted small molecule inhibitor AVA-ADR-001 [38]; the catalytic A-to-I editing inhibitor ADAR1i-124 [39]; Z-DNA-based PROTAC degraders of ADAR1 (Z-PROTACs) [40]; and the ZYS-1 inhibitor [41]— although its ADAR1 selectivity has since been contested [42]. Our findings have translational relevance, as ADAR1 inhibitors may enhance lytic induction therapy in NPC. With EBV lytic-inducing agents already clinically available and multiple ADAR1 inhibitors in development, this strategy may be rapidly adopted into clinical studies. Moreover, the dependence of NPC cells on ADAR1 suggests that ADAR1 inhibition alone may have therapeutic efficacy.

This treatment strategy may be further isolated to EBV-infected cancerous cells by targeting EBV-host interactions that promote ADAR1. For instance, our observation that ADAR1 protein levels are higher in EBV-positive compared to EBV-negative isogenic NPCs may be attributable to EBNA1-mediated translational upregulation of ADAR1 [12] (Fig. 2A). EBNA1 is a well-characterized viral oncogene and several EBNA1 inhibitors have been developed [43, 44]. There may be potential in using EBNA1 inhibitors in combination with lytic reactivation, but further work is required to validate the underlying mechanism. Interestingly, previous reports showed that EBNA1 induces the p150 isoform of ADAR1 [12], while we observed an increase in both p150 and p110 isoforms in the EBV-positive context (Fig. 2A).

Both the p150 and p110 isoforms of ADAR1 edit dsRNA, but their function is context-dependent. ADAR1 p150 can shuttle between the nucleus and cytoplasm, whereas the p110 isoform is believed to stay in the nucleus. However, under stressed conditions, p110 can bind to Exportin-5 upon phosphorylation by MKK6–p38–MSK MAP kinases, thereby promoting its export from the nucleus [45]. Once in the cytoplasm, ADAR1 p110 outcompetes Staufen1 for binding to double stranded structures in the 3’ UTRs of antiapoptotic mRNA [45]. This function prevents their decay and promotes survival of the cell during stress. Although not yet investigated, ADAR1 p110 may be critical in preventing SG formation during stress conditions. Importantly, ADAR1 p110—but not p150—interaction with PKR is induced during poly(I:C) treatment [24]. This rationale agrees with our findings that SGs appear in the absence of ADAR1 and further increase during lytic reactivation (Fig. 5), particularly if p110 is the predominant isoform exploited by EBV in NPC to regulate PKR.

ADAR1 regulates SG formation through multiple mechanisms. For example, Corbet et al. demonstrated that ADAR1 suppresses SG formation through two mechanisms: first, by inhibiting dsRNA-mediated activation of PKR and the ISR; and second, by directly disrupting RNA–RNA interactions that promote SG assembly, independently of its editing activity [46]. Our findings that poly(I:C) induces SGs in the absence of ADAR1 supports a role for the editing-independent function of ADAR1 in preventing SG formation (Fig. 5B). Future work is required to deduce the mechanisms used by ADAR1 to limit SG assembly during EBV lytic reactivation.

Taken together, our study implicates ADAR1-mediated regulation of PKR activity as an essential host mechanism exploited by EBV to evade innate immune surveillance. By upregulating ADAR1, EBV promotes persistent latent infection and oncogenesis in NPC. Inhibiting ADAR1 during forced EBV lytic reactivation represents a promising strategy to enhance lytic induction therapy by rendering EBV vulnerable to ISR activation, thereby impeding viral protein synthesis and replication while inducing cancer cell death.

## Materials and methods

### Cell culture

NPC cell lines were cultured in Roswell Park Memorial Institute (RPMI) 1640 medium (Gibco) supplemented with 10% heat-inactivated fetal bovine serum (FBS) and 1% penicillin-streptomycin (10,000 U/mL; Gibco), and maintained at 37°C in a humidified incubator with 5% CO₂. All NPC cell lines were cultured in the presence of 4 μM Rho-associated protein kinase (ROCK) inhibitor (Y-27632; Enzo), with the exception of HK-1 and C666-1. Human embryonic kidney (HEK)293T cells used for lentivirus production were maintained in high-glucose Dulbecco’s Modified Eagle Medium (DMEM; Gibco) supplemented with 10% FBS and 1% penicillin-streptomycin. Lentiviral transductions were performed in the presence of 8 μg/mL polybrene (Merck). Cells were detached using TrypLE Express Enzyme (1×, no phenol red; Gibco) and passaged before reaching confluence. Cell numbers were determined using a CASY Cell Counter (Roche Innovatis AG) prior to seeding. All cell lines were tested for mycoplasma contamination using the MycoAlert Mycoplasma Detection Kit (Lonza) and authenticated by short tandem repeat (STR) profiling.

### EBV lytic reactivation

Latent EBV in NPC cell lines was induced into lytic reactivation by either transfecting 1 µg/mL of a BZLF1-expressing plasmid or treating cells with a mixture of TPA (20 ng/mL) and NaB (2 mM) for 24–48 hrs at 37°C. Mock transfection or mock treatment was used as a control.

### Incucyte live-cell imaging

Live-cell imaging was performed using an Incucyte SX1 Live-Cell Analysis System. 4000–8000 cells were seeded into 96-well plates and placed into the Incucyte. Phase and green fluorescent images were acquired every 6 hr using a 10x objective lens. The GFP total integrated density (GCU x µm^2^/image) was calculated for each timepoint and plotted as binned averages for each day. The indicated treatments were administered at time 0 hr.

### Lentivirus production

HEK293T cells were seeded to reach 80–90% confluence the following day and transfected in T75 flasks with 3.5 μg pMD2.G, 10.5 μg psPAX2, and 14 μg transfer plasmid using JetPRIME transfection reagent (Sartorius) in Opti-MEM (Gibco). Culture medium was replaced after overnight incubation. Viral supernatants were collected over the subsequent two days, pooled, filtered through a 0.45 μm syringe filter, aliquoted, and stored at −80°C. Transduction efficiency was assessed by measuring BFP-positive cells using flow cytometry.

### MTT assay

Cell viability was assessed using the MTT assay as previously described [47]. Cells were seeded into 96-well plates to reach 20–30% confluence after 24 hr. On days 0, 3, and 5, 20 μL MTT reagent was added to each well and incubated for 4 h at 37°C. The medium was aspirated and formazan crystals were dissolved in 150 μL dimethyl sulfoxide (DMSO). Absorbance at 570 nm and 630 nm was measured after 5–10 min using a Synergy H1 plate reader (BioTek).

### Colony formation assay (CFA)

Colony formation assays were performed as previously described [48]. Cells were seeded at low density (5–10% confluence) in 24-well plates and cultured for 1–2 weeks until colonies became visible. Colonies were fixed in ice-cold methanol for 1 min, stained with crystal violet for 30 min at room temperature in the dark, washed with water, and imaged.

### RNA isolation and cDNA synthesis

Total RNA was isolated using TRIzol (Ambion). Briefly, cell pellets were lysed in 1 mL TRIzol, followed by phase separation with 200 μL chloroform. RNA was precipitated from the aqueous phase using 500 μL isopropanol, washed with 75% ethanol, and resuspended in nuclease-free water. RNA concentration was measured using a NanoDrop ND-1000 spectrophotometer. Complementary DNA (cDNA) was synthesized from 100–1000 ng total RNA using the High-Capacity cDNA Reverse Transcription Kit (Applied Biosystems) according to the manufacturer’s instructions.

### Quantitative Real-Time PCR (qPCR)

cDNA was diluted 1:10 and amplified using the QuantiNova SYBR Green PCR Kit (Qiagen) with gene-specific primers. Each 10 μL reaction contained 2 μL diluted cDNA and 8 μL reaction mix. Plates were centrifuged at 2,000 × g for 2 min before amplification on a QIAquant 96 5plex Real-Time PCR System (Qiagen). Relative gene expression was calculated using the 2^−ΔΔCt method.

### Quantification of EBV copy number

EBV copy number was determined by qPCR targeting EBNA1 using genomic DNA extracted with the QIAamp DNA Blood Mini Kit (Qiagen). A standard curve was generated using the EBV-positive Namalwa cell line, which contains two integrated EBV genomes per cell [49], corresponding to 96–300,000 EBV genome copies.

### Immunoblotting

Whole-cell lysates were prepared in radioimmunoprecipitation assay (RIPA) buffer supplemented with protease and phosphatase inhibitors (Thermo Scientific). Lysates were incubated on ice for 20 min, clarified by centrifugation (14,000 × g, 20 min, 4°C), and protein concentration was determined using the bicinchoninic acid (BCA) assay (Thermo Scientific). Equal amounts of protein (20–30 μg) were mixed with Laemmli sample buffer containing 10% 2-mercaptoethanol, separated by SDS-PAGE, and transferred to polyvinylidene fluoride (PVDF) membranes. Membranes were blocked in 5% milk or bovine serum albumin (BSA) in TBST for 1 h and incubated overnight at 4°C with primary antibodies (Supplementary Table S1), followed by HRP-conjugated secondary antibodies for 1 h at room temperature. Proteins were detected using HRP substrate (Millipore) and visualized with an Amersham Imager 600 (GE Healthcare). Membranes were stripped using 1% sodium hydroxide before re-probing.

### Statistical analysis

Statistical analyses were performed using GraphPad Prism 9. Comparisons between two groups were performed using two-tailed unpaired Student’s t-tests or Welch’s t-tests where unequal variances were detected. Multiple-group comparisons were performed using two-way ANOVA with appropriate post hoc multiple-comparison tests. Data are presented as technical replicates averaged from, or representative of at least two independent biological experiments, unless otherwise stated. Data are presented as mean ± s.e.m., and P < 0.05 was considered statistically significant.

## Statement of translational relevance

Viruses rely on host factors to support their lifecycle, presenting key vulnerabilities for therapeutic intervention. Our study identifies a previously unrecognized dependence of Epstein-Barr virus (EBV) on the RNA-editing enzyme, adenosine deaminase acting on RNA 1 (ADAR1) for both the maintenance of persistent latent infection and successful lytic reactivation in nasopharyngeal carcinoma (NPC). In the absence of ADAR1, lytic induction no longer supports productive viral replication. Instead, viral double-stranded RNA (dsRNA) activates protein kinase R (PKR), leading to eukaryotic initiation factor 2 alpha (eIF2α) phosphorylation, stress granule formation, loss of viral protein expression, and selective death of EBV-positive cancer cells without the production of infectious virions. These findings establish a therapeutic strategy that overcomes a major limitation of conventional lytic induction therapy—the risk of viral dissemination. By combining lytic induction therapy with ADAR1 inhibition, EBV triggers an innate antiviral response, thereby selectively eliminating the virus-infected tumour cells while preventing the generation of viral progeny.

## Abbreviations

ADAR1: Adenosine deaminase acting on RNA 1
dsRNA: Double-stranded RNA
EBV: Epstein-Barr virus
eIF: Eukaryotic initiation factor
GEF: Guanine nucleotide exchange factor
HDAC: Histone deacetylase
IFN: Interferon
IGF2BP3: Insulin-like growth factor 2 mRNA-binding protein 3
ISR: Integrated stress response
KSHV: Kaposi’s sarcoma-associated herpesvirus
MDA5: Melanoma Differentiation-Associated protein
5 NaB: Sodium butyrate
NPC: Nasopharyngeal carcinoma
PACT: PKR-activating protein
PERK: PKR-like endoplasmic reticulum kinase
PKR: Protein kinase R
PRR: Pattern recognition receptor
SG: Stress granule
TPA: 12-O-tetradecanoylphorbol-13-acetate
tRNA: Transfer RNA

## Resource availability

All materials and resources will be made available upon reasonable request from the corresponding author.

## Acknowledgements

We thank Dr. Anna Chi Man Tsang (The Chinese University of Hong Kong) for supplying the HK-1 and NPC43 (EBV positive and negative) cell lines. This work was funded by sponsors of Cancer Research Malaysia. Cancer Research Malaysia is a non-profit research organization, committed to an understanding of cancer prevention, diagnosis and treatment through a fundamental research program. All authors from Cancer Research Malaysia are supported by internal funding contributed by donors and supporters of Cancer Research Malaysia. Schematics were created in BioRender, licence: *Wiebe, S. (2026)* https://BioRender.com/ijztfan.

## Author contributions

Conceptualization, S.W., S.C.C., A.W.Y.C; Investigation, S.W., M.G.N., A.R.; Formal analysis, S.W., M.G.N., A.R.; Data curation, S.W., M.G.N., A.R.; Methodology, S.W., M.G.N., A.R.; Visualization, S.W.; Project administration, S.W., S.C.C., A.W.Y.C.; Funding acquisition, S.C.C., A.W.Y.C; Supervision, S.W., S.C.C., A.W.Y.C.; Writing— original draft, S.W.; Writing—review & editing, All authors.

