## Supplemental information for "Epstein-Barr virus co-opts ADAR1 to evade PKR activation of the integrated stress response in nasopharyngeal carcinoma"

1 Translational Cancer Biology Research Unit, Cancer Research Malaysia, Subang Jaya, Selangor, Malaysia.

2 Department of Oral & Maxillofacial Clinical Sciences, Faculty of Dentistry, Universiti Malaya, Kuala Lumpur, Malaysia.

3 Oral Cancer Research and Coordinating Centre (OCRCC), Faculty of Dentistry, Universiti Malaya, Kuala Lumpur, Malaysia.

### Supplementary Figure S1

Fig. 1A: HK-1 Cas9; sgNT vs. sgA1K and sgA2G; Rep 1 and 2

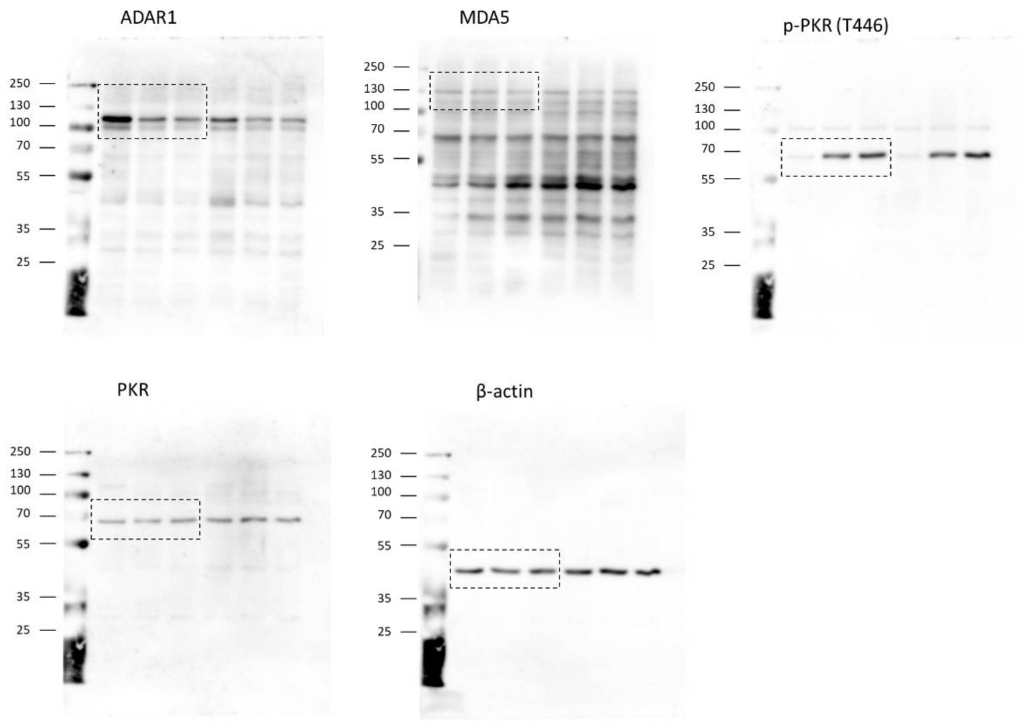

Fig. 1B: HK-1 EBV Cas9; sgNT vs. sgA1K and sgA2G; Rep 1 and 2

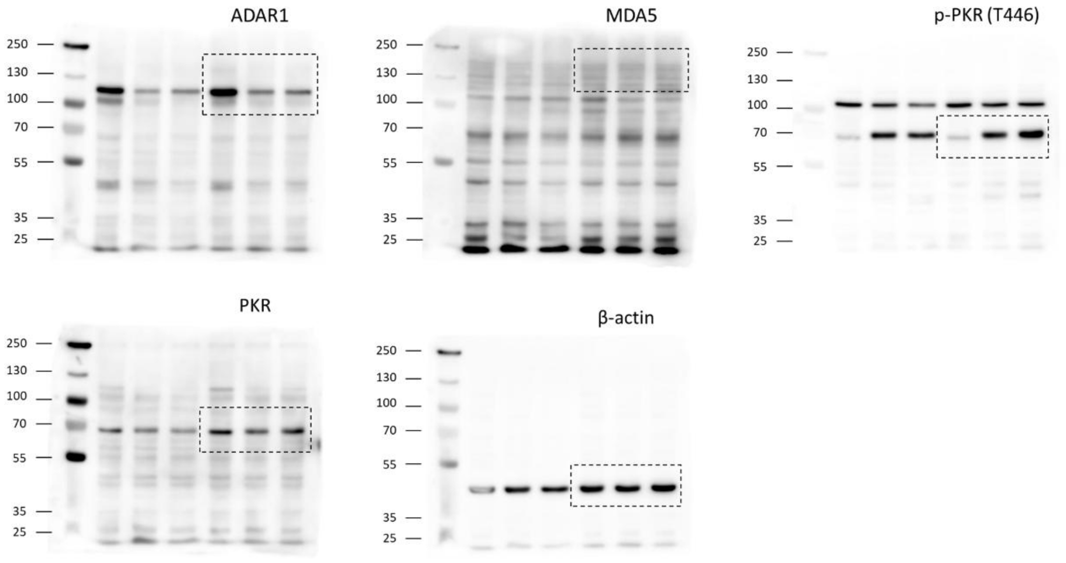

Fig. 1C: C666-1 Cas9; sgNT vs. sgA1K and sgA2G; Rep 1 and 2

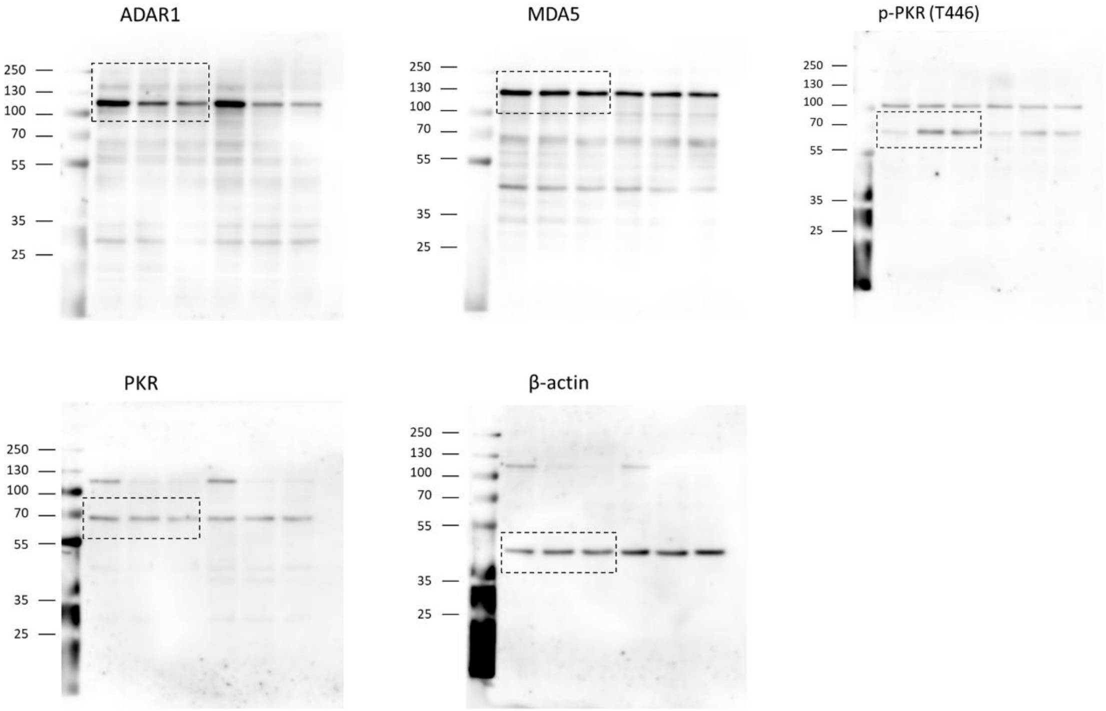

Fig. 1D: NPC268 Cas9; sgNT vs. sgA1K and sgA2G; Rep 1 and 2

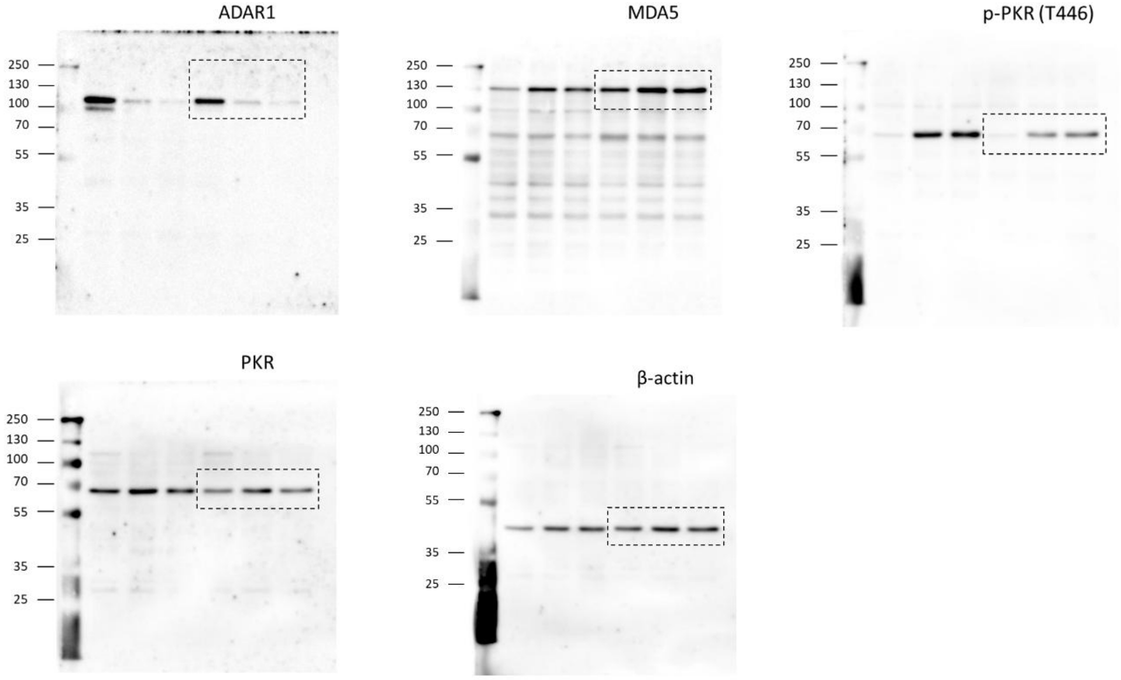

Fig. 2A: NPC43 Cas9; EBV neg vs. pos; sgNT vs. sgA2G

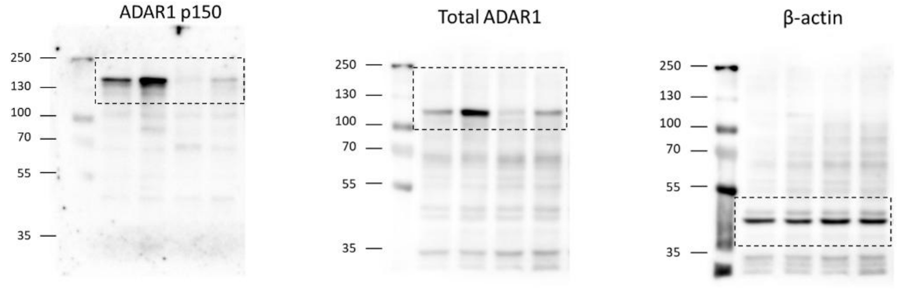

Fig. 2A: HK-1 Cas9; EBV neg vs. pos; sgNT vs. sgA2G

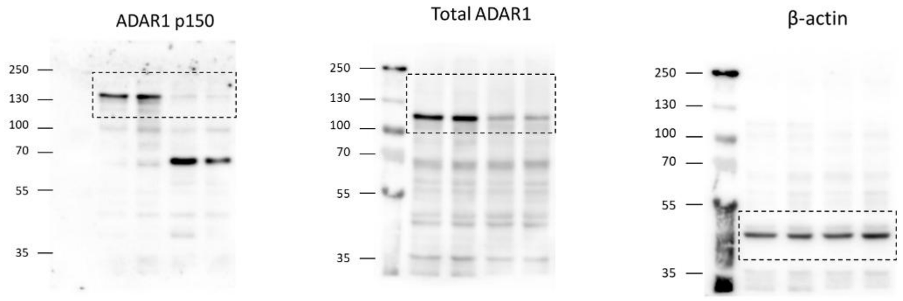

Fig. 4A: HK-1 EBV Cas9; TPA/NaB time course; sgNT vs. sgA2G

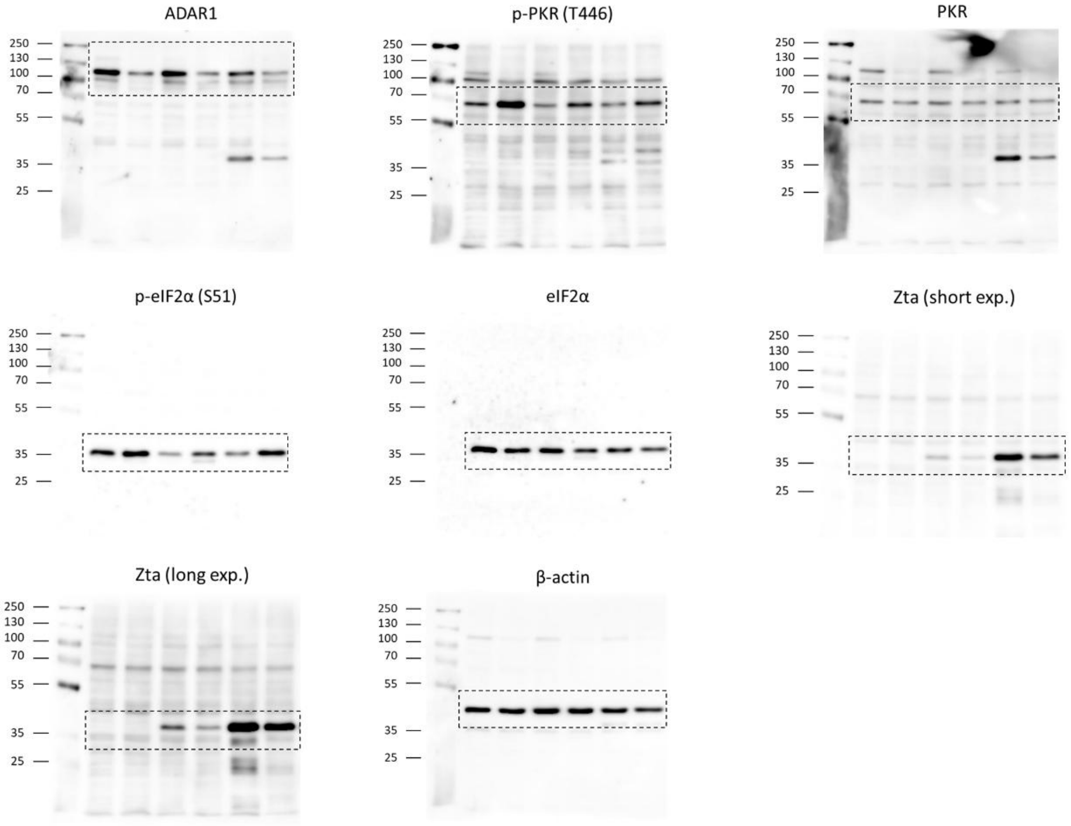

Fig. 4B: NPC268 Cas9; TPA/NaB time course; sgNT vs. sgA2G

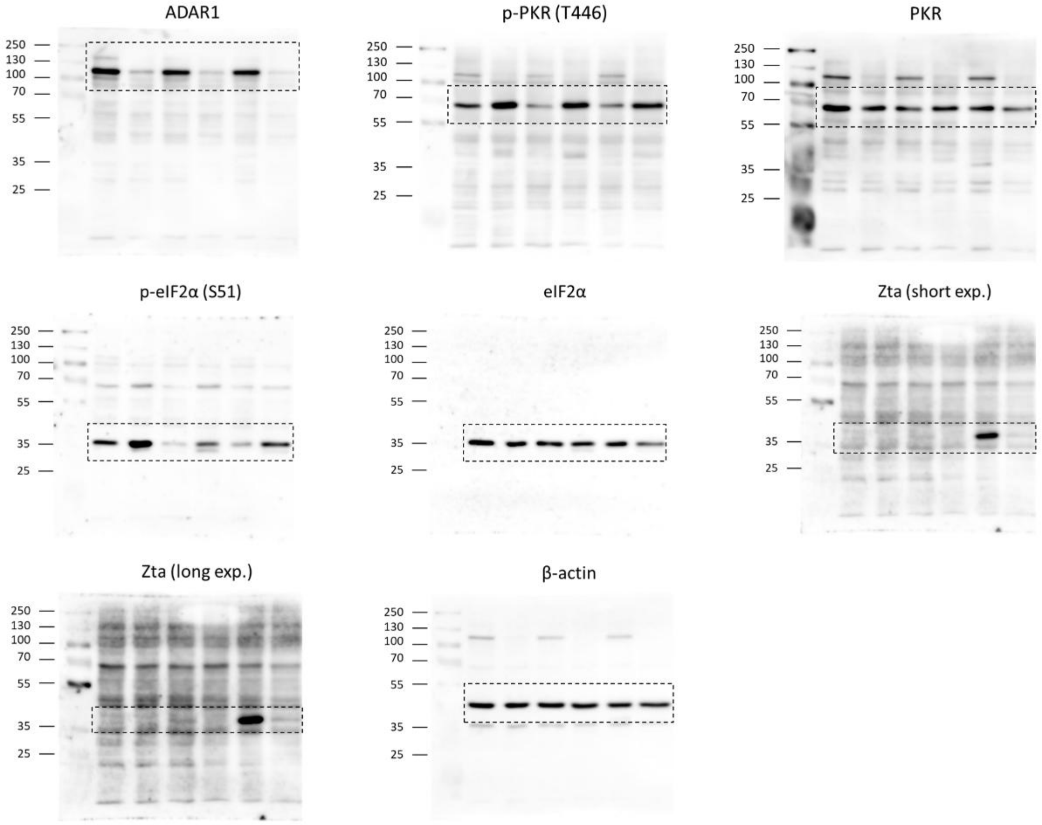

Fig. 4C: HK-1 Cas9; Poly(I:C); sgNT vs. sgA2G vs. sgA2G/PKR

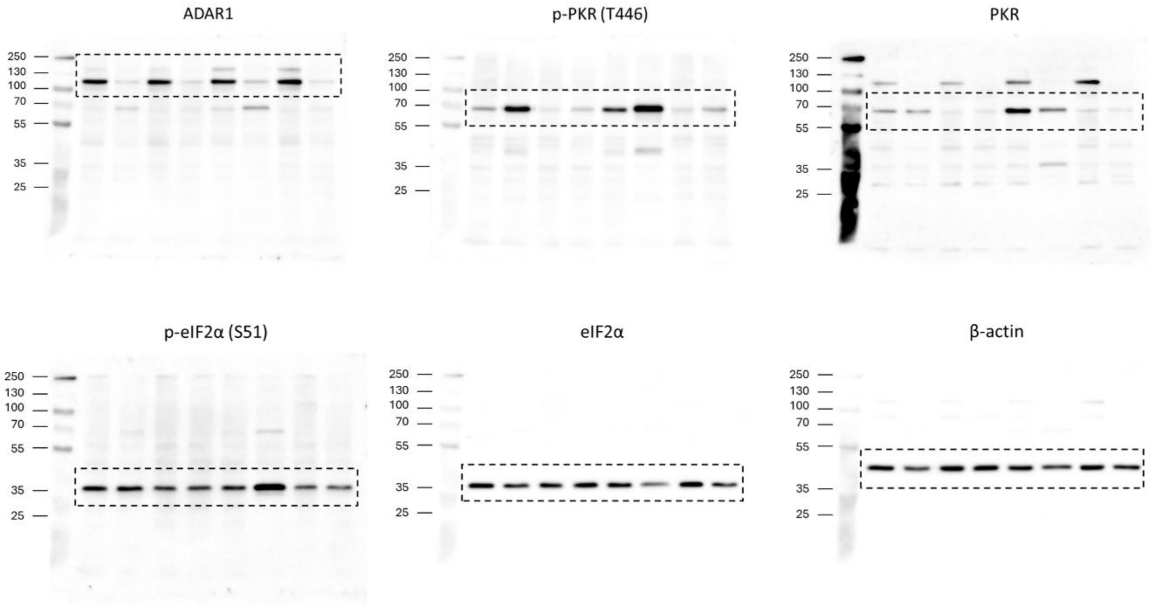

Fig. 4D: NPC43 Cas9; Poly(I:C); sgNT vs. sgA2G vs. sgA2G/PKR

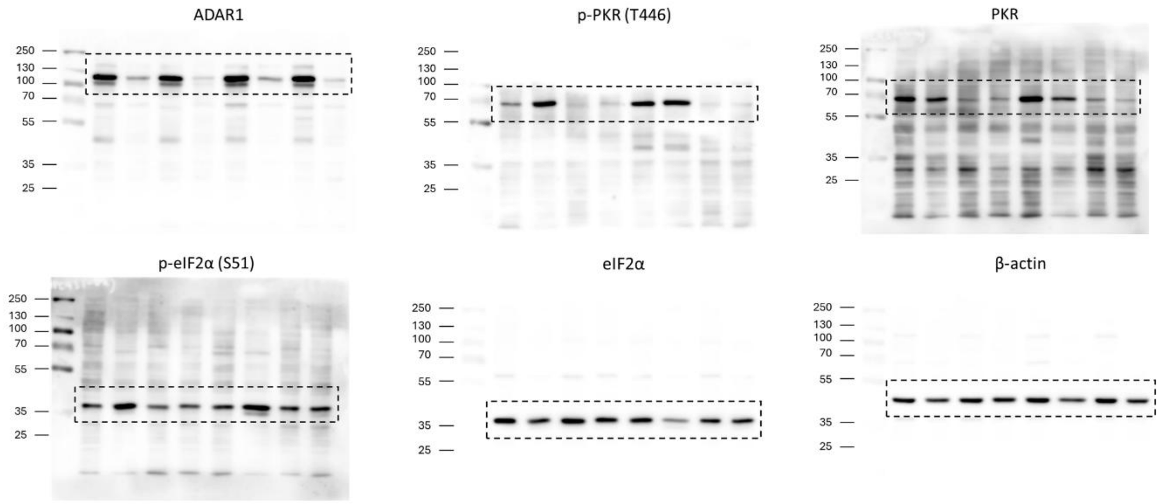

**Figure S1: Uncropped western blots**

Original uncropped western blot images. The dashed box indicates where blots were cropped in the main figures. The protein being probed is indicated above the blot and the molecular weight ladder is located on the left of the blot. Membranes were stripped and re-probed.

**Supplementary Table S1: Key resources**

| Reagent or Resource | Source | Identifier |
|---|---|---|
| **Cell culture** | | |
| RPMI 1640 Medium | Gibco | Cat# 11875093 |
| DMEM, high glucose | Gibco | Cat# 11965092 |
| TrypLE™ Express Enzyme (1X), no phenol red | Gibco | Cat# 12604021 |
| Fetal Bovine Serum | Gibco | Cat# 10270-106 |
| Penicillin-Streptomycin (10,000 U/mL) | Gibco | Cat# 15140122 |
| jetPRIME DNA/siRNA (co-) transfection | Sartorius | Cat# 101000001 |
| Opti-MEM I Reduced Serum Medium | Gibco | Cat# 31985-062 |
| **Antibodies** | | |
| ADAR1 | Santa Cruz Biotechnology | Cat# sc-73408; RRID: AB_2222767 |
| ADAR1 (p150) | Abcam | Cat#: ab126745; RRID: AB_11145661 |
| MDA5 (D74E4) | Cell Signaling Technology | Cat# #5321; RRID: AB_10694490 |
| p-PKR (T446) | Abcam | Cat# ab32036; RRID: AB_777310 |
| PKR (B-10) | Santa Cruz Biotechnology | Cat# sc-6282; RRID: AB_628150 |
| p-eIF2α (S51) | Cell Signaling Technology | Cat# 9721; RRID: AB_330951 |
| eIF2α | Cell Signaling Technology | Cat# 9722; RRID: AB_2230924 |
| EBV ZEBRA (BZ1) | Santa Cruz Biotechnology | Cat# sc-53904; RRID: AB_783257 |
| J2 | Scicons | Cat# 10010500; RRID: AB_2651015 |
| G3BP1 (E9G1M) | Cell Signaling Technology | Cat# 61559S; RRID: AB_2909406 |
| Anti-actin (clone C4) | Millipore | Cat# MAB1501; RRID: AB_2223041 |
| Goat Anti-Rabbit Ig, Human ads-HRP | Southern Biotech | Cat# 4010-05; RRID: AB_2632593 |
| Goat Anti-Mouse Ig, Human ads-HRP | Southern Biotech | Cat# 1010-05; RRID: AB_2728714 |
| Goat anti-human IgG | Southern Biotech | Cat# SB-2040-01 |
| Hoechst 33342 | Thermo Scientific | Cat #62249 |
| Alexa fluor | Invitrogen | Cat# A32728 (mouse); A32732 (rabbit) |

| | | |
|---|---|---|
| **Chemicals** | | |
| Y-27632 Dihydrochloride | Enzo | Cat# ALX-270-333-M005 |
| Puromycin Dihydrochloride | Gibco | Cat# A11138-03 |
| Blasticidin S, Hydrochloride | Merck | Cat# 203350-25mg |
| G-418 Sulfate | Capricorn | Cat# G418-Q |
| MTT reagent | Amresco | Cat# 0793-5g |
| TRIzol | Ambion | Cat# 15596018 |
| QuantiNova SYBR Green PCR Kit (500) | Qiagen | Cat# 208054 |
| Halt Protease and Phosphatase Inhibitor Cocktail | Thermo Scientific | Cat# 1861284 |
| Immobilon Forte Western HRP substrate | Millipore | Cat# WBLUF0500 |
| Ampicillin sodium | TM media | Cat# 69-52-3 |
| Hexadimethrine bromide (polybrene) | Merck | Cat# 28728-55-4 |
| **Commercial Assays** | | |
| High-Capacity cDNA Reverse Transcription Kit | Applied Biosystems | Cat# 4368813 |
| Pierce BCA Protein Assay Kits | Thermo Scientific | Cat# 23227 |
| QIAamp DNA Blood Mini Kit | Qiagen | Cat# 51106 |
| QIAGEN Plasmid Midi Kit | Qiagen | Cat# 12145 |
| QIAquick Gel Extraction Kit | Qiagen | Cat# 28706 |
| **Cell lines** | | |
| HEK293T | | CRL-3216 |
| C666-1 | | CVCL_7949 |
| HK-1 | | CVCL_7084 |
| NPC43 (EBV-negative) | | N/A |
| NPC43 (EBV-positive) | | CVCL_UH64 |
| NPC268 | | Chai et al. 2024 [45] |
| HK-1 EBV | | N/A |
| **Recombinant DNA** | | |
| pKLV2-U6gRNA5(BbsI)-PGKpuro2ABFP-W | Addgene | RRID: Addgene_67974 |
| pKLV2-U6gRNA5(NT)-PGKpuro2ABFP-W | Addgene | RRID: Addgene_163169 |
| pKLV2-U6gRNA5(hPLK1)-PGKpuro2ABFP-W | Addgene | RRID: Addgene_163138 |
| pKLV2-U6gRNA5(hCHAT)-PGKpuro2ABFP-W | Addgene | RRID: Addgene_163147 |
| pKLV2-U6gRNA5(hADAR1_1k)-PGKpuro2ABFP-W | Addgene | RRID: Addgene_208433 |
| pKLV2-U6gRNA5(hADAR1_2g)-PGKpuro2ABFP-W | Addgene | RRID: Addgene_208434 |

| | | |
|---|---|---|
| pKLV2-U6gRNA5(hPKR_2g)-PGKpuro2AmCherry-W | Addgene | RRID: Addgene_208435 |
| **Enzymes** | | |
| T4 Polynucleotide Kinase | NEB | Cat# M0201L |
| T4 DNA ligase | NEB | Cat# M0202L |
| BbsI-HF | NEB | Cat# R3539S |
| **Software and Algorithms** | | |
| Prism 9 | GraphPad | https://www.graphpad.com/features |
| CRISPick | BROAD institute | https://portals.broadinstitute.org/gppx/crispick/public |
| Zen 3.6 | Zeiss | https://www.zeiss.com/microscopy/en/products/software/zeiss-zen.html |

**Supplementary Table S2: CRISPR sgRNA sequences**

| **Gene** | **sgRNA** | **Sequence** |
|---|---|---|
| Non-targeting | sgNT top | CACCGGGAGGTGGCTTTAGGTTTT |
| | sgNT bottom | AAACAAAACCTAAAGCCACCTCCC |
| PLK1 | sgPLK1 top | CACCGTCGGACGCGGACACCAAGG |
| | sgPLK1 bottom | AAACCCTTGGTGTCCGCGTCCGAC |
| CHAT | sgCHAT top | CACCGACGAGCGTTTGCCTCCAAT |
| | sgCHAT bottom | AAACATTGGAGGCAAACGCTCGTC |
| ADAR1 | sgA1K top | CACCGTATTCTAACAGCCCGCTGA |
| | sgA1K bottom | AAACTCAGCGGGCTGTTAGAATAC |
| | sgA2G top | CACCGGATAGCCATGCTGAGCCAC |
| | sgA2G bottom | AAACGTGGCTCAGCATGGCTATCC |
| PKR | sgPKR top | CACCGAAGGCAATACGTACCACTG |
| | sgPKR bottom | AAACCAGTGGTACGTATTGCCTTC |

**Supplementary Table S3: List of primers and probes**

| Name | Primer | Sequence (5′–3′) |
|---|---|---|
| EBNA1 | EBNA1_F | GAGAAGGCCCAAGCACTG |
| | EBNA1_R | CTCCTTGACCACGATGCTTT |
| EBER1/2 | EBER1/2_F | GAGGTTTTGCTAGGGAGGAGA |
| | EBER1/2_R | CACCACCCGGGACTTGTA |
| LMP1 | LMP1_F | GTCCTGTGGGCCATTGTC |
| | LMP1_R | CCCACTCTGCTCTCAAAACC |
| LMP2 | LMP2_F | GACATGAAGAGCACGAAGAGC |
| | LMP2_R | TTCTCATGCTCCTATGGACACTT |
| BZLF1 | BZLF1_F | CCGGCTTGGTTAGTCTGTTG |
| | BZLF1_R | AGCTTATGCAGCACCTCAGC |
| BRLF1 | BRLF1_F | GCTCAGGTCCATCTGTCCAC |
| | BRLF1_R | GGGAGATGGCTGACACTGTT |
| BMRF1 | BMRF1_F | GCGAGGAAAAGGACATCGT |
| | BMRF1_R | CTTCACTTTCTTGGGGTGCT |
| BLLF1 | BLLF1_F | CATCTACAGATTCCAGGCTTACTTG |
| | BLLF1_R | AGCTTCCAATTAACGTCACCA |
| EBNA1 | EBNA-1162F | TCATCATCATCCGGGTCTCC |
| | EBNA-1229R | CCTACAGGGTGGAAAAATGGC |
| | EBNA-1186T probe | (FAM)CGCAGGCCCCCTCCAGGTAGAA(TAMRA) |

| | | |
|---|---|---|
| Beta globin | β-globin-354F | GTGCACCTGACTCCTGAGGAGA |
| | β-globin-455R | CCTTGATACCAACCTGCCCAG |
| | β-globin-402T probe | (FAM)AAGGTGAACGTGGATGAAGTTGGTGG(TAMRA) |
